# Unveiling the links between peptide identification and differential analysis FDR controls by means of a practical introduction to knockoff filters

**DOI:** 10.1101/2021.08.20.454134

**Authors:** Lucas Etourneau, Nelle Varoquaux, Thomas Burger

## Abstract

In proteomic differential analysis, FDR control is often performed through a multiple test correction (*i*.*e*., the adjustment of the original p-values). In this protocol, we apply a recent and alternative method, based on so-called knockoff filters. It shares interesting conceptual similarities with the target-decoy competition procedure, classically used in proteomics for FDR control at peptide identification. To provide practitioners with a unified understanding of FDR control in proteomics, we apply the knockoff procedure on real and simulated quantitative datasets. Leveraging these comparisons, we propose to adapt the knockoff procedure to better fit the specificities of quantitive proteomic data (mainly very few samples). Performances of knockoff procedure are compared with those of the classical Benjamini-Hochberg procedure, hereby shedding a new light on the strengths and weaknesses of target-decoy competition.

## 1 Introduction

Controlling the false discovery rate (FDR) is a well-established practice in most -omic approaches, as it answers a pervasive question: Considering quantitative measurements for many covariates (be they genes, transcripts, metabolites, or proteins) in a set of samples split in at least two different biological conditions, how can we shortlist some differentially expressed ones, while controlling the risk of false positives (*i*.*e*. wrongly selected covariates due to their looking differentially expressed while they are not)? To cope with this, the most commonly used procedure is without a doubt the Benjamini-Hochberg one (BH) [2]. However, due to its large field of application, FDR control has focused a lot of additional efforts in biostatistics, and many authors have proposed to improve upon BH FDR control [3, 8], or have proposed alternative frameworks to do so [1, 6, 20].

In the specific case of proteomics, FDR control is not only used in the aforementioned biomarker selection problem. It is also an essential quality control metric when matching experimental fragmentation spectra onto *in silico* spectra (*i*.*e*., derived from reference database of protein sequences). However, for historical reasons, the associated FDR control is not performed using classical tools from biostatistics. On the contrary, a rather empirical approach termed target-decoy [9] is almost exclusively used. It consists in searching two databases: the first one, referred to as target, containing the genuine protein sequences, and another one, referred to as decoy, containing artefactual sequences. Under the assumption that target mismatches and decoy matches are equally likely, the number of decoy matches can be used to estimate the number of target mismatches, thus opening the door to FDR control.

For a long time, FDR control for peptide identification and for protein differential analysis have been considered as largely independent. However, theoretical connections exist: Notably, it has long been established [16] that if target and decoy databases are searched independently, then the procedure is broadly equivalent to relying on empirical null theory to estimate the FDR in a BH-related way [8]. More recently, it has been shown ([7] that BH procedure could be a user-friendly and computationally attractive alternative to target decoy competition (TDC) (*see* **Note 1**). However, recent developments in theoretical biostatistics have made the links between both approaches to FDR control even tighter. Notably, the authors of [1] have proposed to tackle the biomarker research FDR control using an algorithmic procedure akin to that of TDC. Broadly, this novel approach, denoted as “knockoff-filter,” works as follows. First, knockoff variables are simulated to be as independent as possible from conditions of samples, but yet preserve the covariance structure of the original variables (*see* **Note 2**). Second, a competition is organized between each original variable and its associated knockoff. Third, the proportion of retained knockoffs is used to estimate the proportion of wrongly selected original covariates (see Table 1 for a more detailed comparison with TDC). Conversely, authors have recently leverage the theory underlying knockoff filters to propose improved TDC strategies (see [10]).

**Table 1:**
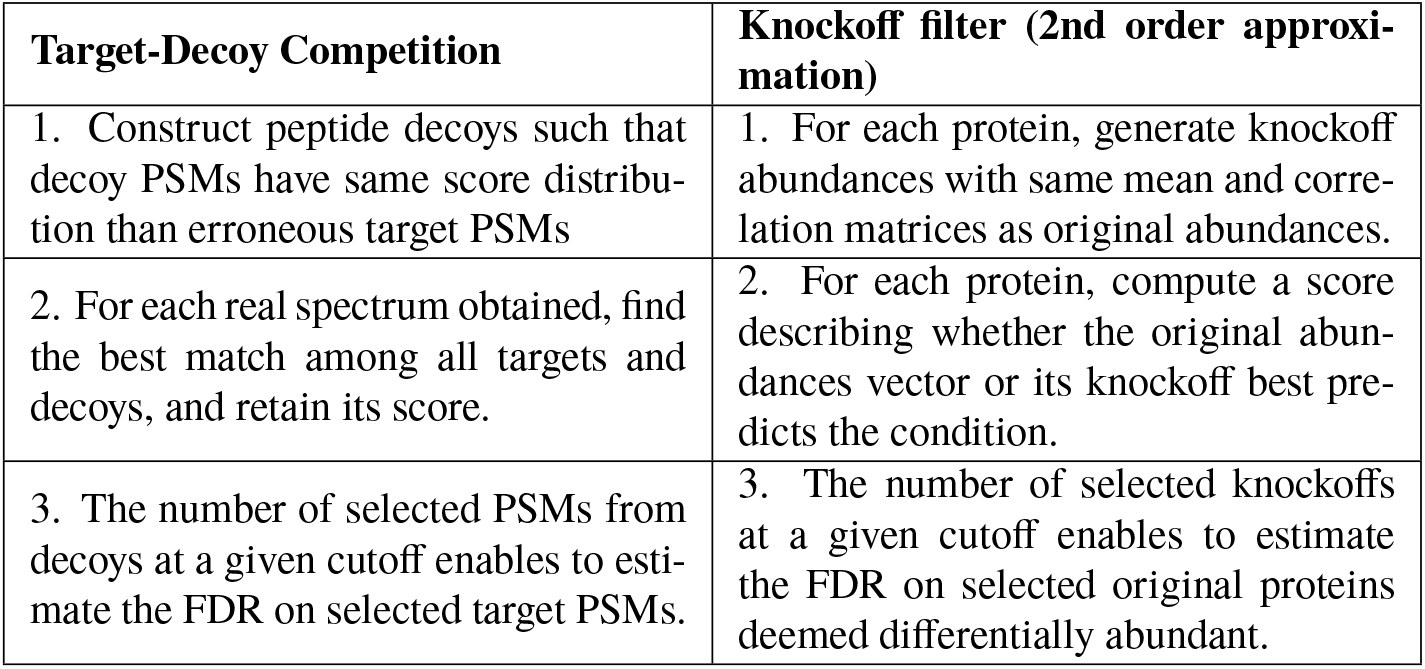
Comparison of the target-decoy and knockoff filter procedures for FDR control. (PSM stands for Peptide-Spectrum Match).

Overall, the framework of knockoff filters is particularly insightful to provide a global understanding of FDR control in proteomics and the purpose of this protocol is to root such unified view on empirical comparisons using both real and simulated data. Interestingly, the results of these comparisons are compliant with empirical knowledge about the various strengths and weaknesses classically associated to each FDR control method.

## 2 Notations

We first start by reviewing commonly used yet conflicting notations in biostatistics and proteomics.

### 2.1 Classical notations in biostatistics

In biostatistics, the false discovery rate (FDR) and the false discovery proportion (FDP) are distinct notions. The FDP corresponds to what was classically and informally referred to as the “true FDR” in proteomics, *i*.*e*., the exact proportion of false positives among the proteins that passed the user-defined selection threshold, and therefore deemed as differentially abundant. Of course, except for benchmark artificial or simulated datasets, this quantity is unknown in practice.

The FDR reads as FDR = 𝔼 [FDP], where 𝔼 stands for the expectation, which broadly amounts to the long run average of the FDP on an infinite number of related experiments subject to stochastic fluctuations. This quantity is also unknown but it is much easier to estimate, and such estimate is classically noted 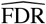. Estimating the FDR is insightful, but unfortunately, not always sufficient [14]. An unbiased FDR estimate is expected to provide a value closed to 𝔼 [FDP]. However, on a given dataset, this value may be larger or smaller than the FDP. While a slightly too large estimate implies a conservative behavior (there will be less false positives than expected among the shortlisted biomarkers), a too small FDR implies a too liberal quality control and subsequent risks in post-proteomics experiments.

To cope with weaknesses of FDR estimation, FDR control procedures have been developed: they rely on more conservative assumptions that yield slightly lesser selected discoveries at a given cut-off parameter. If we note as 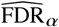 the FDR estimate resulting from controlling the FDR at level *a* (*a* being classically tuned to 1%) it is likely that

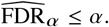

In other words, if one cuts-off a list of putative biomarkers according to an FDR controlled at 1%, the FDR estimate on this very list is likely to be slightly lower than 1%. However, as the FDP remains unknown, it is the only way to safely assume that the FDP is equal to or lower than 1%.

### 2.2 Classical notations in proteomics

In proteomics, most of the notions described above (*see* Subheading 2.1) are conflated. Since the mid-2010s, discriminating between the FDP and the FDR has progressively become standard. However, distinction between FDR (as equal to 𝔼 [FDP], 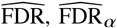, and *a* is scarce. The reason is obvious: except for specific methodological publications, most of them are not useful to the community. Indeed, in practice, a proteomic researcher only needs to manipulate *α*, the cut-off parameter, and to understand that after applying the FDR control accordingly, the FDP is not necessarily strictly equal to *α*, but possibly slightly smaller. However, the everyday language is error-prone: when one says or writes “We selected the putative biomarkers at an FDR of 1%,” what is referred to as FDR is not 𝔼 [FDP], 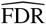, or 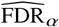, but *a*.

To cope with this, it is possible to rely on other notations. They are not as formal as those of mainstream biostatistics (*see* Subheading 2.1) although they are sometimes reported in mathematics works [4]. However, they are sufficient for a rigorous everyday work in a proteomic lab. Essentially, it amounts to conflate the FDR estimate with *α*, and to define the FDR control as a procedure which provides the following guarantee with a sufficiently high probability:

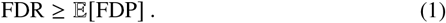

This formulation can be misleading in the sense it gives the impression that the FDR control procedure indeed controls the FDP (*see* **Note 3**). However, it has two advantages: First, it makes the everyday language compliant with the minimum amount of statistical notions possible; second, it simplifies the understanding of other statistical notions such as “q-value” or “adjusted p-value,” as using this formalism, they are simply equivalent to the FDR, as detailed in [5]. In the rest of the protocol, the naming conventions resulting from Eq. 1 are used, so that FDR refers to *α*, the FDR level tuned by the practitioner to perform FDR control.

### 2.3 Other notations used in this protocol

Hereafter, the following mathematical notations are used:

1. *n*: the number of biological samples.
2. *p*: the number of proteins to include in differential analysis.
3. *X* ∈ ℝ^*n*× *p*^ : the matrix of protein abundances, where each row corresponds to a sample and each column corresponds to a protein.
4. *X*_*j*_ : the vector of abundance of the *j* -th protein, *i*.*e*. the *j* -th column of *X*.
5. *x*_*i, j*_ : the abundance value of *j* -th protein for the *i*-th replicate.
6. *y*: the vector representing the condition label (numerical value) of biological samples, of length *n*. For example, the *i*-th coefficient of *y* is 1 if the *i*-th sample comes from the healthy condition, and -1 if it comes from the disease condition.
7. 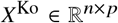 : the knockoff dataset, generated from original dataset matrix *X*.
8. 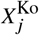: the knockoff vector of abundance of the *j* -th protein.
9. *W* : the vector of scores of all proteins (only the original ones, not the knockoff), of length *p*.
10. *W*_*j*_ : the score associated to the *j* -th protein. A large positive value *W*_*j*_ is evidence that the protein *j* is differentially expressed. It is typically constructed by comparing the predictive power of *X*_*j*_ and 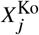of the sample conditions. Swapping *X*_*j*_ and 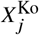 should swap the sign of *W*_*j*_. A null *W*_*j*_ means that both 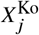 and 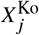 bring the same amount (or lack thereof) of information on the condition.

## 3 Material

### 3.1 R version

R version 4.0.3 (or above) is required to use the following packages. We recommend using an integrated development environment like Rstudio to execute the commands of this protocol. It can be downloaded from https://www.rstudio.com/.

### 3.2 Packages

The following packages are necessary:

1. The packages `knockoff, lars` ([13]), and `glmnet` ([11]) must be installed from the CRAN:

~~~
install. packages (“ knockoff “)
install. packages (“ lars “)
install. packages (“ glmnet “)
~~~
2. `cp4p` [12] provides two datasets with controlled ground truth: They result from analysis of samples containing different abundance of 48 human proteins spiked in a yeast background [19]. The p-values from a Welch *t*-test associated to each protein are also provided, along with functions to apply Benjamini-Hochberg procedure for differential analysis. To install cp4p package, it is first necessary to install the BioConductor [15] packages it depends on:

~~~
if (! requireNamespace (“ Bioc Manager “, quietly = TRUE))
      install. packages (“ Bioc Manager “)
Bioc Manager : : install (“ multtest “)
Bioc Manager : : install (“ limma “)
Bioc Manager : : install (“ q value “)
~~~
3. Then `cp4p` can be installed from the CRAN:

~~~
install. packages (“ cp4p“)
~~~
4. Finally, load the packages in the environment:

~~~
library (cp4p)
library (knockoff)
library (lars)
~~~

### 3.3 Data Format

This protocol relies on a data format which is quite uncommon in proteomics (*see* **Note 4**). The input data *X* on which FDR control is applied should have at least 3 rows, *i*.*e*. at least biological 3 samples in total are needed. The number of proteins to include in differential analysis can be arbitrary. Values of abundance in *X* should be log_2_-scaled.

For conveniency, we use two datasets in this protocol: A dataset resulting from real mass-spectrometry output, called `LFQRatio25` (*see* Subheading 3.4), and a simulated dataset with adjustable parameters (*see* Subheading 3.5).

### 3.4 Data loading from cp4p

The following commands enable to load and prepare `LFQRatio25` dataset [12]:

1. Load the dataset with the following command:

~~~
data (“ LFQRatio25 “)
~~~
2. Then, abundances values for all 6 samples are extracted to form the rows of the `X_yups` variable:

~~~
X_yups = t (LFQRatio25 [, 1 : 6 ])
~~~
3. Similarly, vector `y_yups` contains the condition labels of these samples:

~~~
y_yups = c (1, 1, 1, ™ 1, ™ 1, ™ 1)
~~~
4. For this particular dataset, differentially abundant proteins (or in statistical language, variables under the alternative hypothesis *H*_1_) are known. It is possible to display their name and their index in the list of proteins. These are the 46 first proteins, as the output of this code chunk suggests (*see* **Note 5**):

~~~
mask_human = LFQRatio 25 $Organism == “ human “
names_diff_yups = LFQRatio 25 $Majority. protein. IDs [ mask_human ]
idx _ diff _ yups = which (mask_human)
idx _ diff _ yups
[1] 1 2 3 4 5 6 7 8 9 10 11 12 13 14 15 16 17
[18] 18 19 20 21 22 23 24 25 26 27 28 29 30 31 32 33 34
[35] 35 36 37 38 39 40 41 42 43 44 45 46
~~~
5. Check the dataset to make sure the same dataset is obtained:

~~~
head (X_yups [, 1 : 5 ])
         [,1]     [,2]     [,3]     [,4]     [,5]
A.R1 31.27392 29.48101 29.80982 29.10410 26.85626
A.R2 31.27147 29.46032 29.84163 29.22384 27.11535
A.R3 31.26327 29.45797 29.83771 29.00945 26.94358
B.R1 29.83022 28.04973 28.41002 27.45505 25.71735
B.R2 29.81413 28.02686 28.38101 27.58463 25.74196
B.R3 29.84867 28.00774 28.42514 27.52028 24.62264
~~~

### 3.5 Data simulation

The following commands enable to prepare a simulated dataset:

1. The code below randomly generates a dataset broadly akin to `LFQRatio25`. Due to randomness, it will be different from one run to another. To ensure the results are reproducible and to obtain same results as in the remaining of the protocol, use the following optional command to set the random seed (*see* **Note 6**):

~~~
set. seed (1234)
~~~
2. Tune the parameters of the dataset:

~~~
n_h1 = 50    # Number of proteins differentially abundant
n_rep = 3    # Number of replicates of each condition
p=1500       # Number of proteins
mu = runif (p, 24, 32)
sigma 1 = diag (runif (p, 0, 0.0 2))
sigma 2 = diag (runif (p, 0, 0.02))
mu_diff = c (runif (n_h1, 0.5, 2) * sign (runif (n_h1, ™1, 1)) ,
   rep (0, p–n_h1))
~~~
3. Create and concatenate arrays of both conditions:

~~~
p = length (mu)
X1 = matrix (rnorm (n_rep*p), n_rep) %*% chol (sigma 1)
X2 = matrix (rnorm (n_rep*p), n_rep) %*% chol (sigma 2)
X1 = t (t (X1)+mu+mu_diff / 2)
X2 = t (t (X2)+mu−mu_diff / 2)
X_sim = rbind (X1, X2)
y_sim = c (rep (1, n_rep), rep (−1, n_rep))
idx_ diff_sim = 1 : n_h1
~~~
4. Check the dataset to make sure there are no mistakes:

~~~
head (X_sim [, 1 : 5 ])
         [,1]     [,2]     [,3]     [,4]     [,5]
[1,] 23.96519 28.55181 27.95301 29.64470 31.05108
[2,] 24.04396 28.21652 27.74679 29.56570 31.51248
[3,] 24.05717 28.39634 27.90406 29.74762 31.56869
[4,] 25.65308 29.55612 29.92890 28.21821 30.47228
[5,] 25.74846 29.63377 29.77653 28.30777 30.32441
[6,] 25.89306 29.58248 29.80624 28.33325 30.52107
~~~

## 4 Methods

This section falls into the following subsections:

1. We explain how to apply the original knockoff-filter procedure to control the FDR for differential expression analysis. Precisely, we show how to (1) generate knockoff variables; (2) compute a score for each protein/knockoff pair; (3) select differentially abundant proteins for a predefined target FDR.
2. We detail how to replace the default scoring strategy with other ones, and compare these alternative knockoff procedures to the classical Benjamini-Hochberg (BH) procedure.
3. We propose some code to illustrate the sensitivity of the knockoff filter procedure to the random generation of knockoffs.

### 4.1 Original knockoff procedure

1. Choose the dataset on which applying the knockoff procedure:
  a. To apply it on the `LFQRatio25` dataset, use:

~~~
X_data = X_yups
y_data = y_yups
idx_diff = idx _ diff _ yups
~~~
  b. Alternatively, to apply it on the simulated dataset, use:

~~~
X_data = X_sim
y_data = y_sim
idx_diff = idx_diff_sim
~~~ For the rest of this section, we will use the `LFQRatio25` dataset.
2. Rescale the data to have null mean and unitary variance for each protein abundance vector (*i*.*e*. for each *X*_*j*_) (*see* **Note 7**):

~~~
X_data = scale (X_data)
~~~
3. Execute these commands to generate the knockoff dataset from original data with a fixed seed (*see* **Note 8**):

~~~
set. seed (1234)
X_data_k = create. second order (X_data)
~~~
4. For each protein, compute a score based on the Lasso path of covariates (*see* **Note 9**). An inevitable warning concerning the lack of replicates appears: “one multinomial or binomial class has fewer than 8 observations; dangerous ground.”

~~~
set. seed (1234)
W_lasso = stat. lasso _lambdasmax_bin (X_data, X_data_k, Y_data)
~~~
5. Set the value of targeted FDR, compute the resulting threshold, and select proteins for which their score is above this threshold. The `target_fdr` parameter must be a number between 0 and 1. The `offset` parameter determines which FDR estimator to use, it can be set to either 0 or 1 (*see* **Note 10**). When `offset` is 0, a biased FDR estimate is used, and when `offset` is 1, a non-biased, yet more conservative estimate is used.

~~~
target _ fdr = 0. 05
thres = knockoff. threshold (W_lasso, fdr = target _ fdr, offset = 0)
selected _ lasso = which (W_lasso >= thres)
~~~
6. **This step and the following ones are optional, as they can only be applied for a dataset endowed with a ground truth, such as** LFQRatio25 **or a simulated dataset**. Display the names of proteins selected as differentially abundant at the FDR tuned with the `target_fdr` parameter (here 0.05).

~~~
names _diff_yups [ selected _ lasso ]
[1] P02768upsedyp ALBU_HUMAN_upsedyp - CON P02768-1
[2] O00762upsedyp UBE2C_HUMAN_upsedyp
[3] P00709upsedyp LALBA_HUMAN_upsedyp
[4] P02788upsedyp TRFL_HUMAN_upsedyp
[5] P06396upsedyp GELS_HUMAN_upsedyp
[6] P12081upsedyp SYHC_HUMAN_upsedyp
~~~
7. This code instantiates useful functions to compute the FDP and power from ground truth data. For a certain selection level *α*, the power is defined as

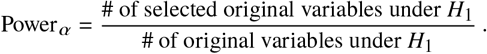

Where *H*_1_ denotes the alternative hypothesis, *i*.*e*. “the protein is differentially abundant.” The power gives a measure of how well our selection covers all the proteins differentially expressed:

~~~
compute_fdp = function (selected, nonzero) {
    if (length (selected) ! = 0) {
         return (1 – sum (nonzero %in% selected) / length (selected))
    }
    return (0)
}
compute_power= function (selected, nonzero) {
    if (length (selected) ! = 0) {
         return (sum (nonzero %in% selected) / length (nonzero))
    }
    return (0)
}
~~~
8. The following code computes the FDP and power of the procedure for a user-defined range of target FDRs (for both offset values):

~~~
FDR = seq (0, 0.5, 0.04)
template = rep (0, length (FDR))
FDP = list (template, template)
POWER = list (template, template)
for (t in 1 : length (FDR)) {
  for (offs in 1 : 2) {
    thres = knockoff. threshold (W_lasso, fdr =FDR[ t ] ,
         offset = offs −1)
    selected = which (W_lasso >= thres)
    FDP [ [ offs ] ] [ t ] = compute_fdp (selected, idx _ diff)
    POWER[ [ offs ] ] [ t ] = compute_power (selected, idx _ diff)
    }
}
~~~
9. Using the results computed at the previous step, the following code displays the FDP and power as a function of the FDR (see Figure 1 for `LFQRatio25` and Figure 2 for simulated dataset):

~~~
par (pty = ‘ s ‘)
cols = c (“ red “, “ blue “, “ black “)
plot (FDR, FDR, type = ‘ l ‘, ylab = “FDP “, xlab = “FDR” ,
     ylim =c (0, 0. 5), xlim =c (0, 0. 5))
lines (FDR, FDP [ [ 1 ] ], col =“ red “)
lines (FDR, FDP [ [ 2 ] ], col =“ blue “)
points (FDR, FDP [ [ 1 ] ], col =“ red “, pch = 1)
points (FDR, FDP [ [ 2 ] ], col =“ blue “, pch = 2)
legend (“ topleft “, legend =c (0, 1, “ y=x “), col = cols ,
    pch=c (1, 2, ™ 1), lty = 1, title =“ Offset “)
plot (1, type =“ n “, ylab = “ Power “, xlab = “FDR” ,
     ylim =c (0, 0. 4), xlim =c (0, 0. 5))
lines (FDR, POWER[ [ 1 ] ], col =“ red “)
lines (FDR, POWER[ [ 2 ] ], col =“ blue “)
points (FDR, POWER[ [ 1 ] ], col =“ red “, pch = 1)
points (FDR, POWER[ [ 2 ] ], col =“ blue “, pch = 2)
legend (“ topleft “, legend =c (0, 1), col = cols, pch=c (1, 2) ,
    lty = 1, title =“ Offset “)
~~~

**Figure 1:**
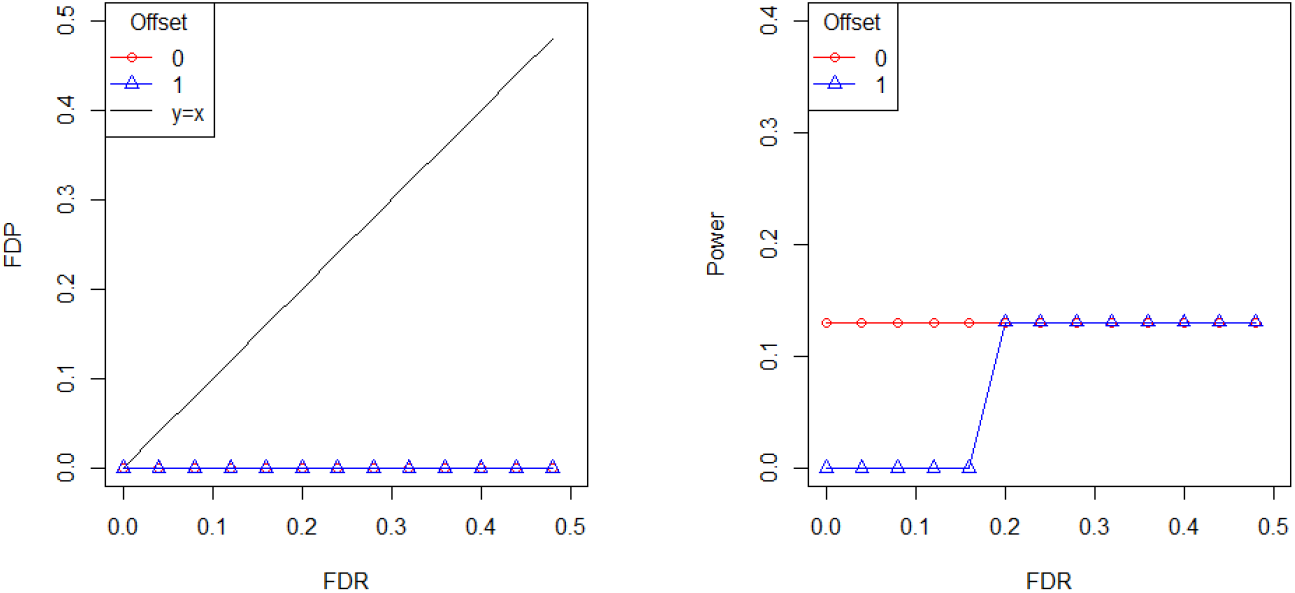
FDP and power *vs*. FDR for `LFQRatio25` dataset, with and without offset, for the knockoff filter procedure with Lasso-based scores.

**Figure 2:**
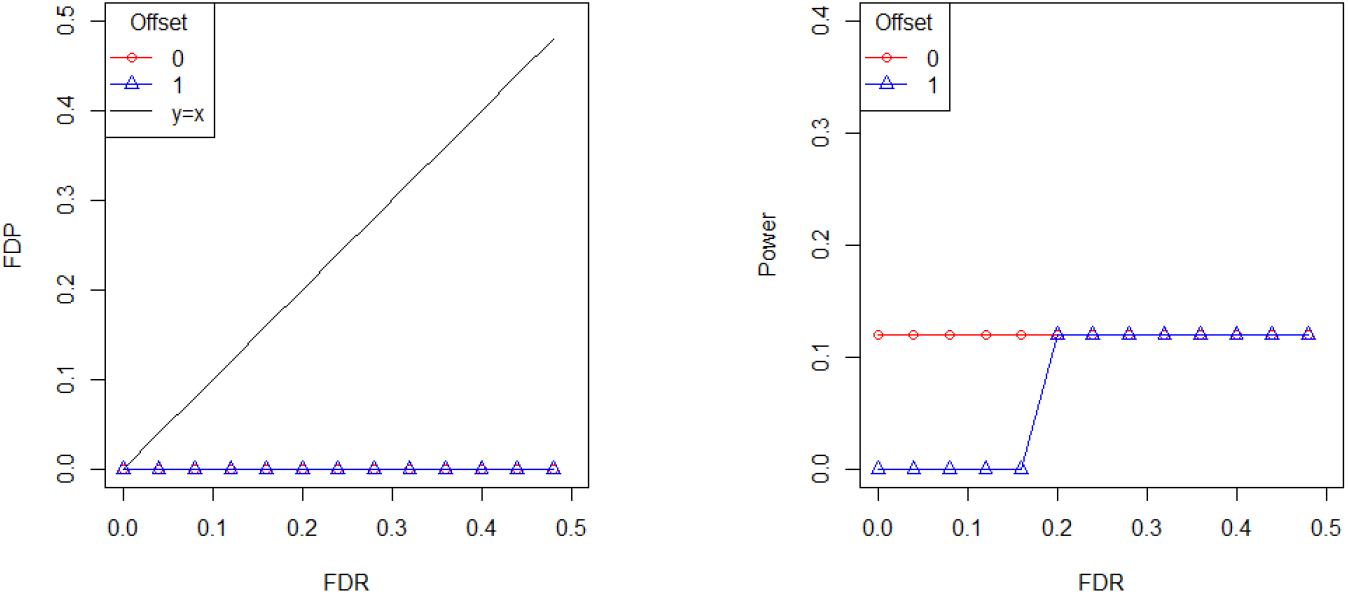
FDP and Power *vs*. target FDR for the simulated dataset, with and without offset, for knockoff procedure with Lasso-based scores.

We notice that FDP and power curves on Figures 1 and 2 are almost always horizontal. This means that variables selected remain the same whatever the FDR target chosen. When the offset equates 1 (unbiased estimator), no proteins are deemed differentially expressed below a certain value of FDR. Thus, even though their are no false positive, there are no true positive either, making the FDR control through knockoff filters practically useless.

We mainly explain this over-conservativeness by the usage of variable selection with the Lasso algorithm, at the step of *W* scores computation. In fact, in the setting *n << p*, the Lasso algorithm will only select *n* variables. This is problematic for differential expression analysis where the total number of samples rarely exceeds the number of *a priori* differentially expressed proteins. On top of that, as very few covariates are selected, and some original variables are much more differentially abundant than all the others, knockoff variables are almost never selected. Thus, estimating the number of false discoveries from the number of selected knockoffs is not appropriate in our cases. These efficiency of variable selection with Lasso is thoroughly discussed in [21].

### 4.2 Scoring methods based on forward stagewise regression and *t*-test

Preliminary experimental comparisons highlighted the knockoff procedure accuracy highly depends on the chosen feature selection algorithm. We herefater describe two procedures that we found to address the issue described above (*see* Subheading 4.1). The first scoring method consists in using forward stagewise selection (FS) algorithm (*see* **Note 11**). The second one is derived from the variable selection procedure classically used in proteomics: it amounts to computing a *t*-test p-value for both original and knockoff variables; then, the final score (*i*.*e*., *W*_*j*_) is defined by the log difference of p-values (LDP) obtained between each original variable and its knockoff.

1. To instantiate the functions that compute the *W*_*i*_ ‘s for the FS and LDP methods, use the following chunks of code (it is advised to run them both, so as to allow subsequent comparisons):
  a. For the FS method:

~~~
stat _ forward _ sel = function (X, X_k, y) {
   Xconcat = c b i n d (X, X_k)
   res = lars (Xconcat, c (1, 1, 1, −1, −1, −1), type =“ for “ ,
       use. Gram = FALSE)
   lambdas = rep (0, 2* ncol (X))
   lambdas [ r e s $ entry ! = 0 ] = res $ lambda [ res $ entry ]
   W_fs = lambdas [ 1 : ncol (X)] −lambdas [ (−1 : n c o l (X)) ]
   W_fs
}
W_fs = stat _ forward _ sel (X_data, X_data_k, y_data)
~~~
  b. For the LDP method:

~~~
stat _ log _ diff _ pval = function (X, X_k) {
      Xconcat = cbind (X, X_k)
      pvals = apply (Xconcat, 2, function (x) { res =
          t. test (x [ 1 : 3 ], x [ 4 : 6 ]) ; return (res $ p. value) })
      pvals _ or = pvals [ 1 : (length (pvals) / 2) ]
      pvals _k = pvals [ (length (pvals) / 2 + 1) : length (pvals) ]
      W_pvals = (−log (pvals _ or) + log (pvals _k))
      W_pvals
}
W_ldp = stat _ log _ diff _ pval (X_data, X_data_k)
~~~
2. Plot the histogram of *W*_*i*_ ‘s to better visualize the selection process (see Figure 3 for `LFQRatio25` dataset):

~~~
hist (W_ldp [ W_ldp ! = 0 ], col =c (rep (“ red “, 2), rep (“ grey “, 4) ,
     rep (“ blue “, 1 1)), main =“ Histogram of W”, xlab =“W”)
axis (1, at =c (™ 5, ™2, 0, 2, 5, 1 0))
~~~
3. To illustrate the interest of using FS and LDP within the knockoff filter procedure, we compare those two approaches with the classically used Benjamini-Hochberg (BH) procedure. Depending on the dataset being `LFQRatio25` or the simulated one, the code differs:
  a. With `LFQRatio25`, the p-values resulting from Welch *t*-test are provided in the dataset:

~~~
pvals = LFQRatio25 [, 7 ]
res = adjust. p (pvals, pi0. method = 1)
~~~
  b. With the simulated dataset, p-values must be computed beforehand (a Welch *t*-test is also used here):

~~~
pvals = apply (X_data, 2, function (x) } res = t. test (
    x [ 1 : n_rep ], x [ (n_rep + 1) : (2 * n_rep) ]) ;
    return (res $ p. value) })
res = adjust. p (pvals, pi0. method = 1)
~~~
4. Compute the FDP and power for BH and knockoff filter procedure with LDP and FS methods (with offset=1), at different FDR levels:

~~~
FDP = list (template, template, template)
POWER = list (template, template, template)
W _list = list (W_fs, W_ldp)
for (t in 1 : length (FDR)) {
  for (W_idx in 1 : 2) {
    thres = knockoff. threshold (W _list [ [ W_idx ] ], fdr =FDR[ t ], offset = 1)
    selected = which (W _list [ [ W_idx ] ] >= thres)
    FDP [ [ W_idx ] ] [ t ] = compute_fdp (selected, idx _ diff)
    POWER[ [ W_idx ] ] [ t ] = compute_power (selected, idx _ diff)
  }
  selected _ bh = which (res $ adjp $ adjusted. p<=FDR[ t ])
  FDP [ [ 3 ] ] [ t ] = compute_fdp (selected _b h, idx _ diff)
  POWER[ [ 3 ] ] [ t ] = compute_power (selected _b h, idx _ diff)
}
~~~
5. Finally plot the FDP and power *vs*. FDR level, as illustrated on Figures 4 and 5, respectively for the `LFQRatio25` and simulated datasets):

~~~
par (pty = ‘ s ‘)
cols = c (“ red “, “ blue “, “ orange “)
leg = c (“ Knockoff w F. S. “, “ Knockoff w log diff. “, “B-H. “)
plot (FDR, FDR, type = ‘ l ‘, ylab = “FDP “, xlab = “FDR” ,
     ylim =c (0, 0. 6), xlim =c (0, 0. 1 5))
for (i i n 1 : 3) }
  lines (FDR, FDP [ [ i ] ], col = cols [ i ])
  points (FDR, FDP [ [ i ] ], col = cols [ i ], pch= i)
}
legend (“ topleft “, legend = leg, col = cols, pch = 1 : 3 ,
    title =“ Procedure “)
plot (1, type =“ n “, ylab = “ Power “, xlab = “FDR” ,
     ylim =c (0, 1. 2), xlim =c (0, 0. 1 5))
for (i i n 1 : 3) }
  lines (FDR, POWER[ [ i ] ], col = cols [ i ])
  points (FDR, POWER[ [ i ] ], col = cols [ i ], pch= i)
}
legend (“ topleft “, legend = leg, col = cols, pch = 1 : 2 ,
    title =“ Procedure “)
~~~

**Figure 3:**
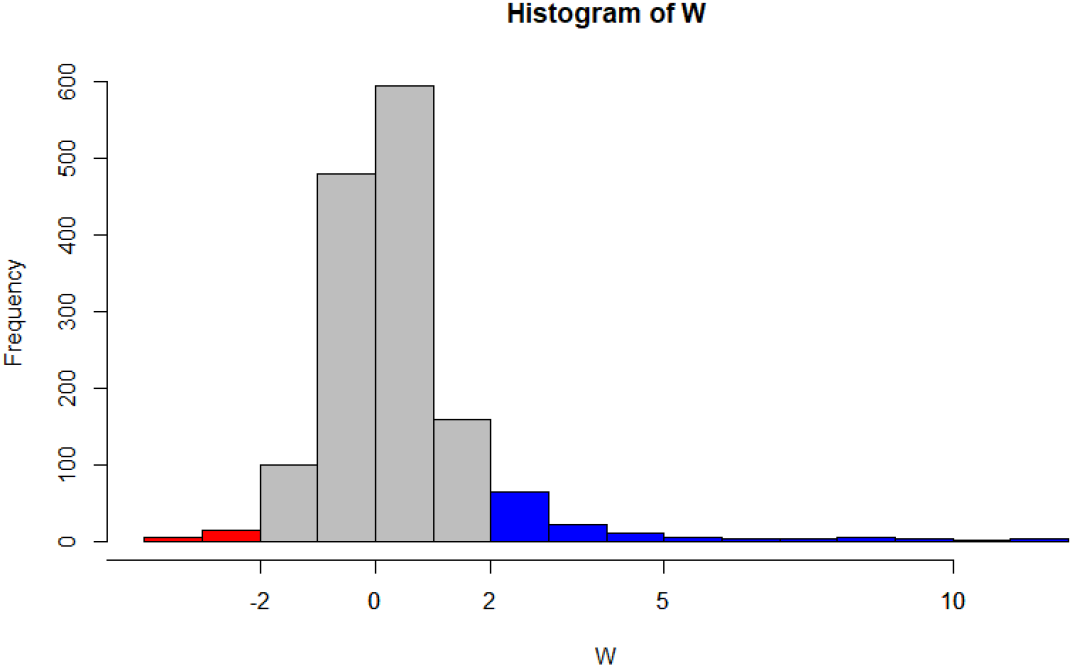
Histogram of scores *W*_*i*_’s obtained with log diff of p-values scoring method, on LFQRatio25 dataset. The blue area correspond to original variables that are selected, and the red area represent knockoff variables selected, both at a threshold of 2 (hence, a conservative FDR estimate at a selection threshold of 2 reads 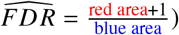.

**Figure 4:**
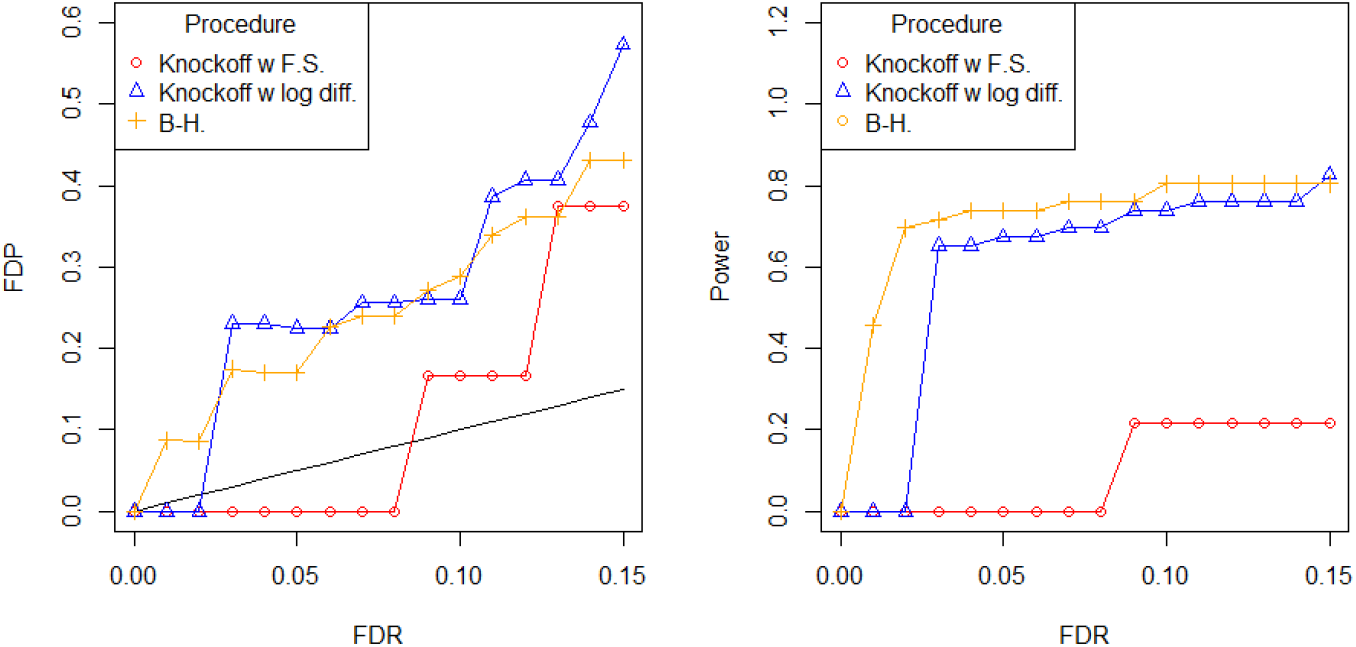
FDP and power *vs*. target FDR for knockoff filter procedure with offset=1 applied with forward stagewise selection and log diff of p-values scoring, and Benjamini-Hochberg procedure, obtained with `LFQRatio25`.

**Figure 5:**
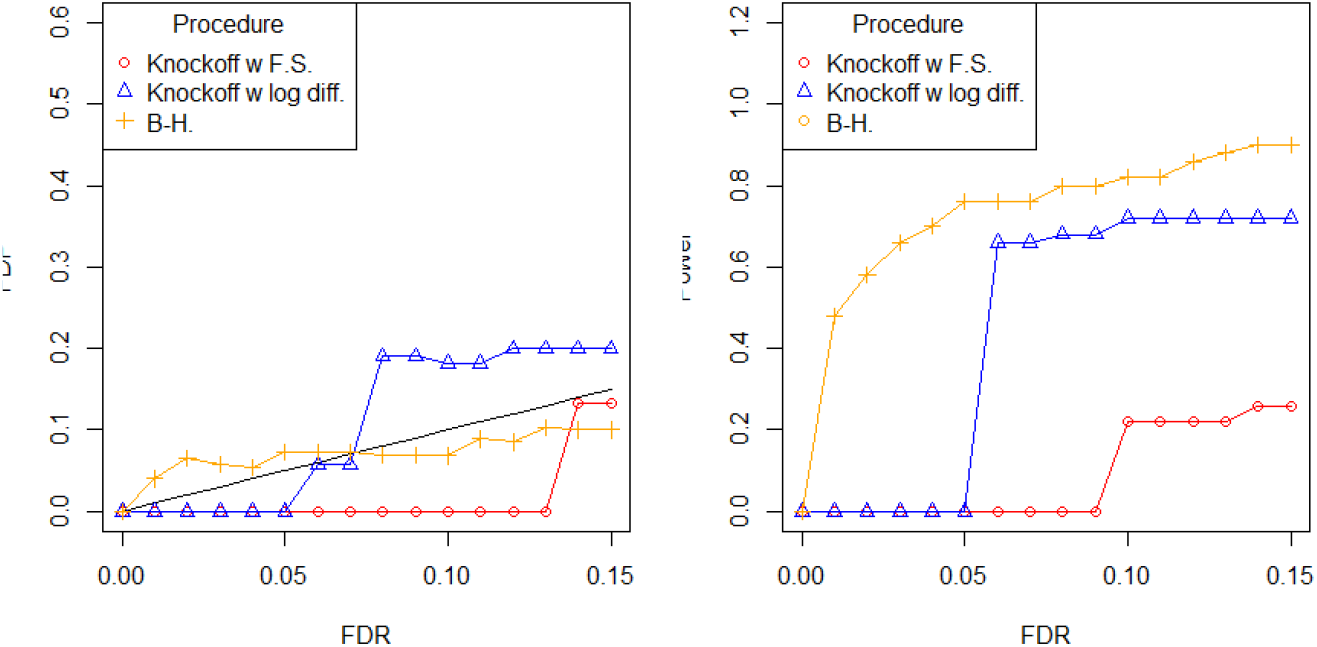
FDP and power *vs*. target FDR for knockoff procedure with offset=1 applied with forward stagewise selection and log diff of p-values scoring, and Benjamini-Hochberg procedure, obtained with simulated data.

We observe that the knockoff filter procedure with LDP broadly follows the same trend as the BH one on `LFQRatio25` (see Figure 4). By construction, the LDP scores is never null, yielding a rather symmetric distribution (see Figure 3). The largest positive scores (depicted in the right hand tail) result from differentially abundant proteins, while the left hand one amounts to selected knockoff proteins. The distribution being more symmetric than when using the Lasso, it is possible to select a larger subset of proteins at a given FDR. However, when using the FS based scores, knockoff filters roughly behaves as with the Lasso, yielding a greater but yet insufficient power.

Finally, the BH procedure also yields anti-conservative results on `LFQRatio25`, as the FDP is always higher than the FDR. However, this can be explained by other preprocessing steps (match between runs, normalization, imputation, etc.) which tends to shrink the within-condition variance prior to differential analysis as well as to increase the risk of false positives that are not accounted by FDR control. Indeed, Benjamini-Hochberg is conservative on simulated data (see Figure 5).

### 4.3 Sensitivity of FDR control to knockoff used

Knockoff generation with `create.second_order` function (*see* Subheading 4.1, **Step 3**) involves the random draw of a knockoff matrix (similarly to the random generation of decoy sequences). Hence, on a given dataset, running two consecutive FDR control procedures with knockoff filters should lead to slightly different results. We hereafter propose several experiments to illustrate the sensitivity of the knockoff filter procedure to the knockoff generation, as well as to evaluate its magnitude.

1. Generate 30 knockoff datasets and store them in a list (depending on the machine, this step may last between 30 minutes to an hour):

~~~
set. seed (3456)
n_k = 30
l_k = list ()
for (i in 1 : n_k) {
  l_k [ [ i ] ] = create. second _order (X_data)
}
~~~
2. Apply the knockoff filter procedure to each knockoff series, with FDR varying from 1% to 15%. In this example, the scoring method used is LDP. For all the knockoff series, the effective FDP *vs*. FDR curves are iteratively plotted, leading to a display akin to that of Figure 6. The proteins selected at an FDR of 5% for each knockoff series are retained in a matrix referred to as scores:

~~~
par (pty = ‘ s ‘)
FDR = seq (0.01, 0.15, 0.01)
FDP < POWER < matrix (rep (0, n_k* length (FDR)), nrow=n_k)
scores = matrix (rep (0, n_k* n c o l (X_sim)), nrow=n_k)
plot (FDR, FDR, type = ‘ l ‘, ylab = “FDP “, xlab = “FDR” ,
     ylim =c (0, 0. 7), xlim =c (0, 0. 1 5))
for (i in 1 : n_k) {
 W = stat _ log _ diff _ pval (X_data, l _k [ [ i ] ])
  for (t in 1 : length (FDR)) }
    thres = knockoff. threshold (W, f d r =FDR[ t ], offset = 1)
    selected = which (W >= t h r e s)
    FDP [ i, t ] = compute_fdp (selected, i d x _ diff)
    if (FDR[ t ] == 0. 0 5) {
      scores [ i, selected ] = 1
    }
  }
  lines (FDR, FDP [ i, ], col = i)
}
legend (“ topleft “, legend = “ y=x “, lty = 1, col =“ black “)
~~~
3. Finally, plot a heatmap featuring the scores matrix which highlights with different colors the selected proteins under *H*_0_ and *H*_1_ for each knockoff filter series, at an FDR target of 5%. (see Figure 7):

~~~
par (mar=c (5, 5, 2, 8), xpd=TRUE, mgp=c (1, 1, 0))
heights = sort (col Sums (scores), decreasing = T ,
    index. return = T)
heights _ in _ plot = heights $ix [ heights $ x > 0 ]
submat = scores [, heights _ in _ plot ]
submat [ (submat == 1) & t (matrix (r e p (heights _ in _ plot > 46 ,
    nrow (scores)), ncol =nrow (scores))) ] = 2
image (t (submat), col =c (“ grey “, “ blue “, “ red “) ,
    xlab =“ Proteins (selected atleast once) “, a xe s =F)
mtext (text =c (paste (“ Knockoff “, c (1, 1 5, 3 0))), side = 2, line = 0. 1 ,
    at = seq (0. 0, 1, 1 / 2), l a s = 1, cex = 0. 9)
legend (“ topright “, inset =c (−0. 23, 0) ,
    legend =c (“ Selected H_0 “, “ Selected H_1 “, “ Not selected “) ,
    fill =c (“ red “, “ cyan “, “ grey “))
~~~

Figures 5 and 7 emphasize the important variability resulting from the random nature of knockoff filters. To counter this variability, [18] proposes a method to aggregate multiple knockoffs. In fact, similar sensitivity has already been commented upon with target-decoy procedures [17], so it seems to be a problem ubiquitous to FDR control procedures which involve simulating artifactual data under the null hypothesis. Finally these observations provide an intuitive support for the tools described in [10], which relies on multiple decoy databases to construct a knockoff-like score.

**Figure 6:**
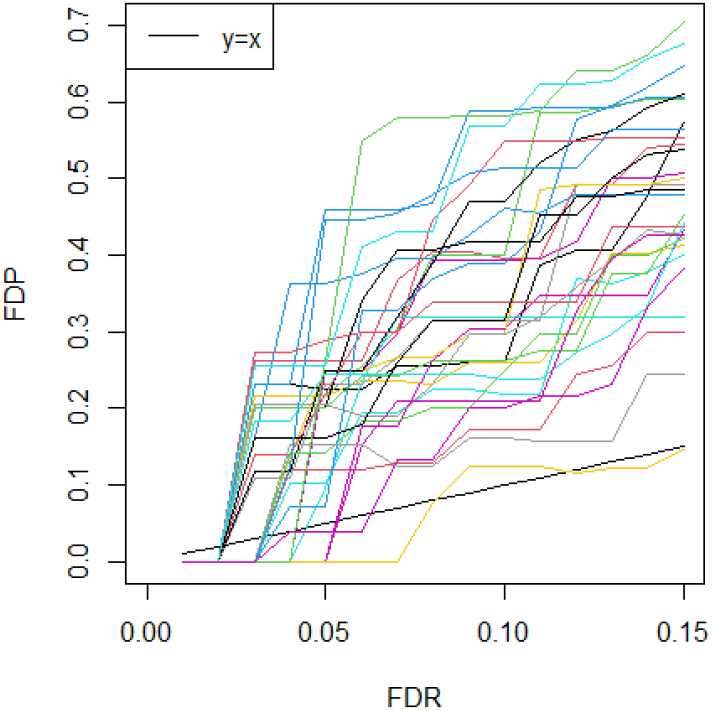
Curves of FDP *vs*. FDR for 30 different Knockoff procedure, applied with log diff of -values score on `LFQRatio25` dataset.

**Figure 7:**
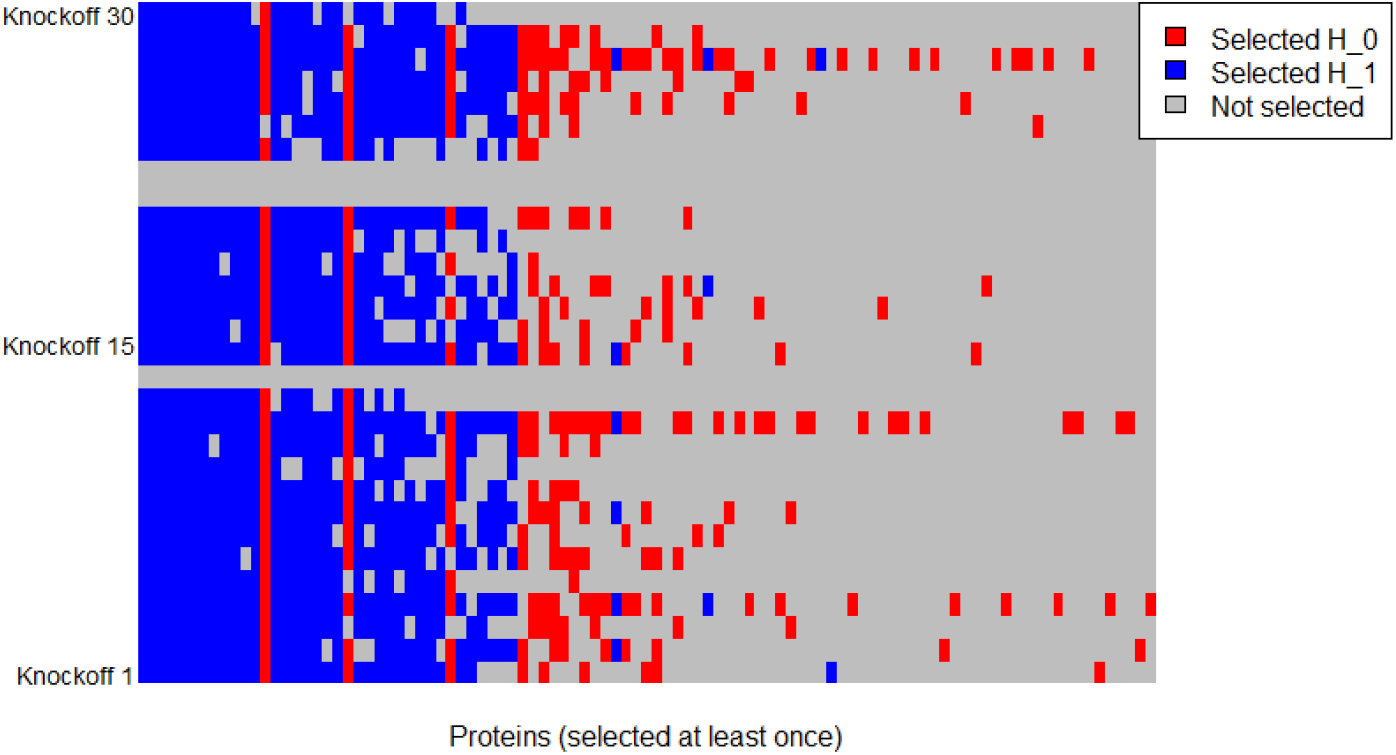
Proteins selected according to 30 different knockoff procedure iterations (using LDP score) on the LFQRatio25 dataset. Blue cells depict original differentially abundant proteins (human proteins) that were selected using a given knockoff. Similarly, red cells depict non-differentially abundant proteins (yeast proteins) mistakenly selected. Proteins are sorted from the most selected one (left hand side) to the least selected one (right hand side).

